# Compound 21, a two-edged sword with both DREADD-selective and off-target outcomes in rats

**DOI:** 10.1101/2020.05.01.072181

**Authors:** Raphaël Goutaudier, Véronique Coizet, Carole Carcenac, Sebastien Carnicella

## Abstract

Although Designer Receptors Exclusively Activated by Designer Drugs (DREADDs) represent a technical revolution in integrative neuroscience, the first ligands used were not as selective as expected. Compound 21 (C21) was recently proposed as an alternative, but *in vivo* characterization of its properties is not sufficient yet. Here, we evaluated its potency to selectively modulate the activity of nigral dopaminergic (DA) neurons through the canonical DREADD receptor hM4Di using TH-Cre rats. In males, 1 mg.kg^−1^ of C21 strongly increased nigral neurons activity in control animals, indicative of a significant off-target effect. Reducing the dose to 0.5 mg.kg^−1^ circumvented this aspecific effect, while activated the inhibitory DREADDs and selectively reduced nigral neurons firing. In females, 0.5 mg.kg^−1^ of C21 induced a transient and residual off-target effect that may mitigated the inhibitory DREADDs-mediated effect. This study raises up the necessity to test selectivity and efficacy of chosen ligands for each new experimental condition.

## Introduction

Designer Receptors Exclusively Activated by Designer Drugs (DREADDs) are chemogenetic tools that represent one of the major breakthroughs of the last ten years in integrative neuroscience (Roth, 2016; Campbell and Marchant, 2018). Combining the precision of genetics with pharmacology, DREADDs provide a remote, prolonged and reversible control of neuronal or extra-neuronal subpopulations via conditional expression and allow the study of complex phenomena in awake animals. As such, they were elegantly used to easily induce tonic modulation, affording an alternative to optogenetics which is more adapted for phasic modulation (Goutaudier et al., 2019), and to study the implication of different neural system in various behaviors such as feeding, memory, pain or motivation (reviewed in Whissell et al., 2016). Initially described as a “lock and key” system, DREADDs are G-protein-coupled receptors that rely on the combination of a mutated muscarinic receptors, that have lost their affinity for acetylcholine, and a designed drug which binds to the mutated receptor with potentially otherwise no pharmacological activity (Armbruster et al., 2007). Two Designed Receptors were originally and are commonly used for DREADD modulation: hM3Dq, coupled to Gq protein which increases neuronal activity and hM4Di, coupled to Gi protein which decreases neuronal activity. A next generation of DREADDs deriving from other endogenous metabotropic and ionotropic receptors were developed over time (Aldrin-Kirk and Björklund, 2019). Similarly, different DREADDs ligands have been developed. The first DREADD ligand was Clozapine-N-oxide (CNO), a derived metabolite of the atypical antipsychotic clozapine, described to be devoid of endogenous activity at moderate doses (Armbruster et al., 2007) and was widely used to activated DREADDs during the past decade. However, the selectivity of this compound was challenged by different studies, where it was observed that CNO induced behavioral off-target effects in both mice and rats which did not express DREADDs (Baerentzen et al., 2019; MacLaren et al., 2016). In addition, Gomez et al. (2017) reported that CNO was not the real DREADDs activator since it was not able to cross the blood brain barrier (BBB) and was in fact back-metabolized in low doses of clozapine. However, behavioral investigations quickly showed that even low doses of clozapine induce anxiety-related behaviors in naïve animals (Ilg et al., 2018; Mahler and Aston-Jones, 2018), indicating that this molecule is not appropriated as a DREADDs ligand (see also Goutaudier et al., 2019). All these observations together led to the necessity of developing new ligands. As such, a second generation of ligands was engineered leading to the creation of three new synthetic ligands: compound 21 (C21) developed by Chen et al. (2015), JHU37152 and JHU37160 (JHUs) developed by (Bonaventura et al., 2019). Compared to JHUs, C21 has been developed earlier and as such, has gained interest in the field. For instance, Thompson et al., 2018 have demonstrated that C21 from 0.3 to 3 mg.kg^−1^ was sufficient to activate DREADDs in mice and to induce selective behavioral alterations. C21 appears therefore to be an interesting DREADDs activator. However, it remains to be fully characterized in other species and other experimental conditions, as caution is needed since designed ligands could have different outcomes depending on the doses, species, strains or gender used (Goutaudier et al., 2019). In the present study, we aimed at further document the *in vivo* properties of C21 by extending its DREADDs application to transgenic TH-Cre rats. Indeed, TH-Cre rats are frequently used in combination to DREADDs as they appear as a powerful tool for the investigation of tonic modulation of mesolimbic and nigrostriatal dopaminergic (DA) systems in motivated, cognitive and affective behaviors (Beloate et al., 2016; Boekhoudt et al., 2016; Mahler et al., 2014; Runegaard et al., 2019). However, no one has tested yet the potential efficiency and selectivity of C21 in this experimental model.

## Results

We thereby infused into the substantia nigra pars compacta (SNc) of male and female TH-Cre rats, a floxed virus encoding for the inhibitory DREADDs hM4Di coupled to mCherry (*♂-hM4Di* and *♀-hM4Di*). Meanwhile, a floxed virus encoding only for mCherry was infused in control groups (*♂-mCherry* and *♀-mCherry*) (**Figure 1**). Before assessing the potential effects of C21 on SNc neuronal activity by using extracellular electrophysiology (**Figure 2A, Figure S1),** we first verified that basal firing rates were comparable between our different experimental groups, since DREADDs may have a constitutive activity (Saloman et al., 2016). We found neither differences between *♂-hM4Di* and *♂-mCherry* animals (23.6 Hz ± 1.9 and 21.2 Hz ± 2.9 respectively, t(39) = 0.65, p = 0.521) nor between *♀-hM4Di* and *♀-mCherry* animals (38.8 Hz ± 7.3 and 26.7 Hz ± 4.3 respectively, t(13) = 1.48, p = 0.164). We also verified that basal neuronal activity and recording remain stable over time in groups of mCherry and hM4Di animals only treated with saline (no effect of time whatever the transgene condition: F_s_ < 0.98, p > 0.46, partial η^2^ < 0.12; data not shown; n = 8). Then, we first assessed the effect of 1 mg.kg^−1^ of C21 in *♂-mCherry* animals (**Figure 2B, D**). We observed a robust and persistent increase in the firing rate of nigral neurons, indicating that, even within the recommended range of doses (Thompson et al., 2018), C21 can have strong non-DREADDs mediated pharmacological effects. Importantly, decreasing the dose of C21 to 0.5 mg.kg^−1^ allowed to completely circumvent this aspecific effect on SNc neuronal activity (main effect of treatment: F_(1, 13)_ = 16.08, p < 0.01; of the time: F_(8, 98)_ = 6.38, p < 0.001; and treatment × time interaction: F_(8, 98)_ = 5.44, p < 0.01; **Figure 2D**, *left panel*). Overall, a 100%-increase was observed between 90 to 180 minutes after the injection of C21 at 1 mg.kg^−1^ compared to vehicle, an effect that was absent at 0.5 mg.kg^−1^ (main effect of the treatment: F_(1, 13)_ = 18.38, p < 0.001, partial η^2^ = 0.59; of the transgene: F_(1, 13)_ = 17.42, p < 0.01, partial η^2^ = 0.59; and treatment x time interaction: F_(1, 13)_ = 15,78, p < 0.01, partial η^2^ = 0.55; **Figure 2D**, *middle and right* panel). We next tested whether the dose of 0.5 mg.kg^−1^ of C21, devoid of off-target effect in the SNc, was sufficient to activate the DREADDs in *♂-hM4Di* animals and to produce significant *in vivo* chemogenetic effects. As shown on **Figure 2C and E**, 0.5 mg.kg^−1^ of C21 induced in *♂-hM4Di* but not in *♂-mCherry* animals, a significant reduction of SNc neuronal firing rate, as expected from the activation of an inhibitory receptor selectively expressed on DA neurons. This decrease became evident 90 minutes after the injection of C21 and ended 120 minutes later (main effect of transgene: F_(1,17)_ = 5.89, p < 0.05; marginal effect of time F_(8,130)_ = 1.97, p = 0.055 and no significant transgene x time interaction F_(8,130)_ = 1.67, p = 0.113; **Figure 2E**, *left panel*). Overall, a 30%-decrease was observed between 90 to 180 minutes after injection of 0.5 mg.kg^−1^ of C21 compared to vehicle and the mCherry control condition (main effect of the treatment: F_(1, 17)_ = 6.8, p < 0.05, partial η^2^ = 0.29; of the transgene: F_(1, 17)_ = 7.46, p < 0.05, partial η^2^ = 0.38; and treatment x time interaction: F_(1, 17)_ = 13.28, p < 0.01, partial η^2^ = 0.44; **Figure 2E**, *middle and right panel*). Notably, we also observed a complete recovery of the basal firing rate in *♂-hM4Di* rats at 240 minutes post-injection. This indicates that, consistently with the DREADDs approach, this effect was reversible, and not due to a loss of signals along time. These finding indicate that, in male TH-Cre rats, 0.5 mg.kg^−1^ of C21 is sufficient to potently activate hM4Di in TH-Cre rats, without inducing endogenous off-target effects.

**Figure 1.**
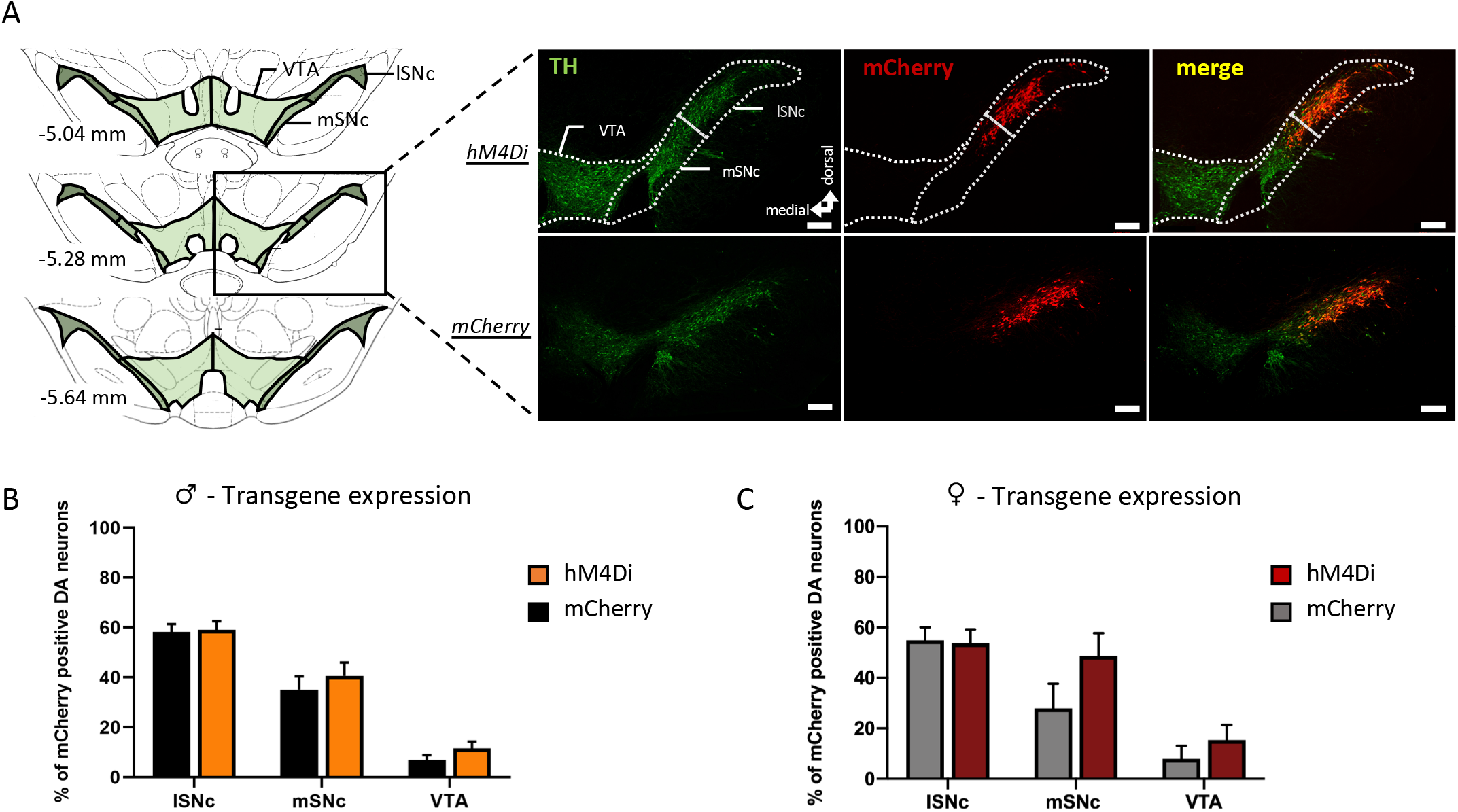
hM4Di-mCherry and mCherry expression in mesencephalic DA neurons of TH-Cre rats. **(A)** *On the left*, schema of the three levels of mesencephalon used for quantified viral expression with three areas: lateral SNc (ISNc), medial SNc (mSNc) and Ventral Tegmental Area (VTA). The black rectangle indicates the level at which representative images, *on the right*, were taken to illustrate TH immunostaining and hM4Di-mCherry or mCherry expression. **(B-C)** Percent of transgenes expression in the lSNc, mSNC and VTA for males **(B)** (hM4Di, orange, n = 18 hemispheres, 13 animals; mCherry, black, n = 23 hemispheres, 16 animals) or females **(C)** (hM4Di, red, n = 7 hemispheres, 7 animals; mCherry, grey, n = 8 hemispheres, 6 animals). A similar pattern of expression was observed between males and females with a gradient of expression from lSNc to the VTA (F_s_ > 18.83, p < 0.001, partial η^2^ > 0.49), whatever the transgene conditions (F_s_ < 2.45, p > 0.16, partial η^2^ < 0.06). For each area, the given percent of expression correspond to the mean expression of the three levels of mesencephalon. Scale bar: 200 μm. Data were expressed as the mean number of quantified hemispheres for which recordings were included +/- SEM

**Figure 2.**
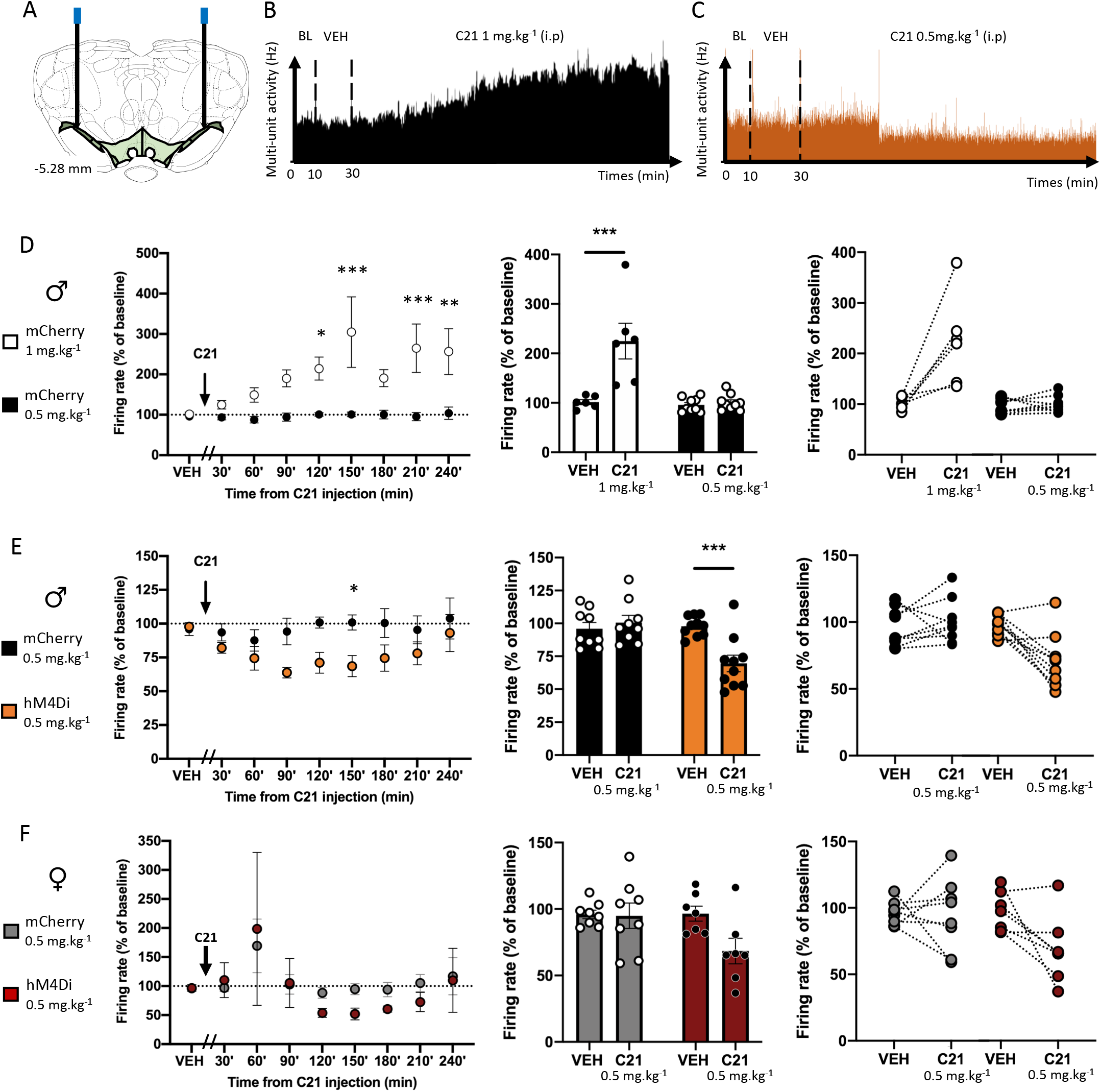
Dose dependent effect of C21 on the activity of SNc neurons expressing hM4Di-mCherry or mCherry. **(A)** Schema of the bilateral electrodes implantations. **(B)** Representative data obtained during recording from male mCherry rat treated with 1 mg.kg^−1^ of C21. **(C)** Representative data obtained during recording from male hM4Di rat treated with 0.5 mg.kg^−1^ of C21. **(D - F)** *On the left*, effect of C21 along time on SNc neuronal firing rate, during vehicle (VEH, a 20-minutes interval) and C21 periods (30-minutes intervals), normalized to 10-minutes baseline recording. *In the middle and on the right*, mean neuronal firing rate, during the VEH period and between the 90 and 180 minutes post-C21 injection intervals, normalized to baseline. **(D)** Effect of C21 in mCherry male rats treated with 1 (white, n = 6 recordings, 4 animals) or 0.5 (black, n = 9 recordings, 7 animals) mg.kg^−1^. **(E)** Effect of C21 in mCherry (black, n = 8 recordings, 7 animals) or hM4Di (orange, n = 10 recordings, 8 animals) male rats treated with 0.5 mg. kg^−1^. **(F)** Effect of C21 in mCherry (grey, n = 8 recordings, 6 animals) or hM4Di (red, n = 7 recordings, 7 animals) female rats treated with 0.5 mg.kg^−1^. Data were expressed as the mean number of recording sides +/- SEM. BL: baseline, VEH: vehicle. *P < 0.05, **P < 0.01, ***P< 0.001.

As brain responses may differ between males and females (Choleris et al., 2018), we next investigated the effect of 0.5 mg.kg^−1^ of C21 in female TH-Cre rats. As in male, this dose of C21 selectively decreased the firing rate of nigral neurons *in♀-hM4Di* rats (**Figure 2F**). This effect was however not detected as significant (**Figure 2F**, *left panel:* no significant effect of the transgene: F_(1, 13)_ = 0.28, p = 0.605; neither of time: F_(8, 95)_ = 1.88, p = 0.072; nor time x transgene interaction: F_(8, 95)_ = 0.23, p = 0.977; **Figure 2F**, *middle and right panel*: marginal effect of treatment: *F_(1,13)_* = 3.63, p = 0.079, no significant effect of the transgene: F_(1, 13)_ = 3.13, p = 0.1, or significant treatment x transgene interaction: F_(1, 13)_ = 3.13, p = 0.1), consistent with lower effect sizes of DREADDs as compared to male (**Figure 2F**, *middle and right panel:* partial η^2^ = 0.22, partial η^2^ = 0.18, partial η^2^ = 0.19, for the treatment, the transgene an treatment x transgene interaction respectively). This is probably due to the fact that, in contrast to male, a residual aspecific effect of C21, highlighted by a clear transient increase of 169% of the firing rate at 60 min post-injection (Fig 2F), likely mitigated the following DREADDs-mediated effect. (**Figure 2F**, *left panel*).

## Discussion

We demonstrated here for the first time that C21 possesses both specific and aspecific effect on rats depending on doses used. In males, at 0.5 mg.kg^−1^, C21 activated hM4Di with a potent *in vivo* effect, without inducing off-target effect. This led to a reversible inhibition of nigral neurons activity selectively in hM4Di-expressing animals. Conversely, at 1 mg.kg^−1^, C21 induced a robust and long-lasting increase of SNc neurons activity in hM4Di-lacking animals. In females, this aspecific effect was also transiently observed, in both hM4Di-expressing and hM4Di-lacking animal, with the dose of 0.5 mg.kg^−1^, meaning that precaution must be taken in studies using both genders. This is critical because scientists working on transgenic lines often used males and females to obtain larger cohorts (e.g., (Bonaventura et al., 2019; Gomez et al., 2017; Saloman et al., 2016; Xia et al., 2017). Relative potent affinity of C21 for some endogenous receptors may account for the off-target effect evidenced in the present study. Indeed, C21 may exhibit similar affinity for serotoninergic 5-HT2 Gi-coupled and histaminergic H1 Gq-coupled receptors than for hM4Di behaving potentially as a competitive antagonist of these receptors (Goutaudier et al., 2019; Jendryka et al., 2019; Thompson et al., 2018). Given that 5-HT2 Gi-coupled receptors are expressed on SNc DA neurons (Cornea-Hébert et al., 1999; Fink and Göthert, 2007), by blocking this inhibitory receptor, C21 can promote SNc neurons activity (Di Giovanni et al., 1999). In addition, blocking H1 Gq-coupled receptors that are located on nigral GABAergic neurons can lead to a reduction of the GABAergic inhibition on nigral DA neurons and therefore enhance their activity (Dringenberg et al., 1998; Korotkova et al., 2002). Although these two hypotheses remain speculative and deserve further investigations, it appears not unlikely that, depending on the dose, C21 exhibits such off-target effect on SNc neuronal activity.

Finally, this study demonstrates for the first time that C21 can be a potent DREADDs activator in rats. It also clearly illustrates that, because DREADDs derive from endogenous receptors and rely on the use of pharmacological compounds, they are unlikely to be fully devoid of off-target effects, even if new ligands are proposed each year and help to maximize this approach. These effects will always depend on the dose, the species, the strains and the gender used. Therefore, regardless of the chosen ligand, a “model-dependent” approach must be adopted to assess the selectivity and efficiency of the ligand for every new experimental condition prior any behavioral experiment.

## Materials and methods

### Animals

29 males and 13 females TH-Cre rats (breeding at the Plateforme Haute Technologie Animal, La Tronche) were included in this study. They were housed in a 12h/12h reverse light cycle, with food and water *ad libitum*. At the beginning of the experiments, the males weighed between 240 and 430g and females weighed between 200g and 320g. All experimental protocols complied with the European Union 2010 Animal Welfare Act and the new French directive 2010/63, and were approved by the French national ethics committee no. 004.

### Stereotaxic viral infusion

Animals were anesthetized with a mixed intraperitoneal (i.p.) injection of ketamine (Chlorkétam, 60 mg.kg^−1^, Mérial SAS, Lyon, France) and xylazine (Rompun, 10 mg.kg^−1^, Bayer Santé, Puteaux, France). Then local anesthesia was provided by a subcutaneous injection of lidocaine (Lurocaine, 8 mg. kg^−1^, Laboratoire Vetoquinol S.A., France) on the skull surface and animal were secured in a Kopf stereotaxic frame under a microbiological safety post (PSM). Coordinates for SNc injections were determined according to Paxinos and Watson, 2007, adjusted to the body weight and set at, relative to bregma: −4.3 mm (AP), ±2.4 mm (ML), −7.9 mm (DV). Animals were infused bilaterally with 1 μl of AAV5-hSyn-DIO-hM4Di-mCherry (10^12^ particles.ml^−1^, Addgene, Watertown, Massachusetts, États-Unis, #44362-AAV5) or 1 μl of AAV5-hSyn-DIO-mCherry (10^12^ particles.ml^−1^, Addgene, #50459-AAV5). The virus was infused at a rate of 0.2 μL.min^−1^ using microinjection cannula (33-gauge, Plastic One, USA) connected to a 10 μL Hamilton syringe and a microinjection pump (Stoelting Co., Wood Dale, IL). After injection, the cannula remained *in situ* for 5 min before withdrawal to allow the injected solution to be absorbed into the parenchyma. The skin was sutured, disinfected, and the animal placed in a heated wake-up cage, before being replaced in its home-cage after complete awakening and monitored for a couple of days.

### Reagent

C21 (Hello Bio, Bristol, UK) was dissolved in 0.9% saline and kept at −20°C before testing. All the injections were given intraperitoneally, at 0.5 or 1 mg.kg^−1^ (at a volume of 1 mL.kg^−1^). A vehicle solution (NaCl 0.9%) was prepared and kept in the same conditions.

### *In vivo* extracellular electrophysiology

At least two weeks after viral infusion, we performed extracellular multiunit recordings to assess neuronal activity of the SNc. Rats were anesthetized continuously with isoflurane and body temperature was maintained at 37°C with a thermostatically controlled heating blanket. Two tungsten electrodes (Phymep, Paris, France) were implanted bilaterally into the SNc using the coordinates determined according to Paxinos and Watson, 2007 and set at: −4.3 mm (AP, bregma), ±2.3 mm (ML, bregma) and −6.5 mm (DV, brain surface). Coordinates between infusion and electrophysiology were slightly changed to avoid the area of mechanical injury induced by the injection. Extracellular voltage excursions were amplified, band-pass filtered (300 Hz–10 kHz), digitized at 10 kHz and recorded directly onto computer disc using a Micro 1401 data acquisition system (Cambridge Electronic Design [CED] Systems, Cambridge, UK) running CED data capture software (Spike 2). Once electrodes were implanted and signal stabilized, baseline (BL) without treatment was recorded during 10 minutes before i.p. administration of a vehicle solution (VEH - NaCl 0.9%). The VEH period of 20 minutes was followed by i.p. administration of C21 (1 mg.kg^−1^ or 0.5 mg.kg^−1^), for a recording period of 240 minutes. Then, the position of SNc recording sites were marked with a small lesion caused by passing 10 μA DC current for 1 min through the tungsten recording electrode. Firing rate activity was normalized by the baseline firing rate activity of the first 10 minutes. Recordings with more than 25% of variation of the firing rate activity between the baseline pre-injection and the vehicle periods were excluded from the study. One recording per hemisphere were performed. Recordings were excluded after histological and immunohistological analyses (see below) when the recording site was outside the SNc and/or when DREADDs or control virus expression in the SNc was absent.

### Tissue Preparation and histological validation

At the end of the experiment, rats were deeply anesthetized by isoflurane saturation and transcardially perfused with 0.9% NaCl (100 mL) followed by 4% paraformaldehyde (300 mL, PFA) in phosphate-buffered saline (PBS). After decapitation, brains were extracted and post-fixed for 24h in 4% PFA. They were then cryoprotected in 20% sucrose/PB for 24h and frozen in isopentane cooled to −50°C on dry ice. Coronal sections (30 μm) of mesencephalon were cut using a cryostat (Microm HM 525; Microm, Francheville, France). Placement of electrodes were verified by Cresyl violet staining and visualized with the ICS FrameWork computerized image analysis system (TRIBVN, 2.9.2 version, Châtillon, France), coupled to a light microscope (Nikon, Eclipse 80i) and a Pike F-421C camera (ALLIED Vision Technologies, Stadtroda, Germany) for digitalization. Meanwhile, floating coronal section of three levels of the mesencephalon were selected according to (Drui et al., 2014) for assessment of DREADDs expression.

### TH-immunohistochemistry and DREADDs expression localization

To assess DREADDs expression in DA mesencephalic regions, immunostaining for tyrosine hydroxylase (TH) was performed. Free-floating 30 μm thick coronal sections were washed with TBS and incubated for 1 h in 0.3% Triton X-100 in TBS (TBST) and 3% normal goat serum (NGS). They were then incubated with primary monoclonal mouse anti-TH antibody (mouse monoclonal MAB5280, Millipore, France, 1/2500) diluted in TBST containing 1% NGS overnight (4°C). Then, slices were incubated with a green fluorescent conjugated goat anti-mouse Alexa 488 antibody (1/500, Invitrogen™, Waltham, Massachusetts, USA) for 1h30 at room temperature. They were finally mounted on superfrost glass slides, with Aqua-Poly/Mount (Polysciences, Inc., Germany). Fluorescent pictures of TH labelling and mCherry expression were taken using a slide scanner (Z1 Axioscan, Zeiss Göttingen, Germany), at x20 magnification and analyzed with ImageJ. DREADD expression was quantified for each hemisphere by comparing the number of TH-labeled-mCherry-positive neurons with the number of TH-labeled neurons within three areas: the lateral SNc (lSNc), the medial SNc (mSNc) and the Ventral Tegmental Area (VTA). We have also verified first that mCherry was only detected in TH-positive neurons. Fluorescent illustrations presented in this article were taken with a laser-scanning confocal microscope (LSM710, Zeiss,). Z-stacks of digital images were captured using ZEN software (Zeiss).

### Data analyses

For DREADDs expression, data were expressed as the mean number of quantified hemispheres for which recordings were included +/- SEM (number of quantified hemispheres and animals for each group are detailed in figure legends). For extracellular electrophysiology, data were expressed as the mean number of recordings +/- SEM (number of included recordings and animals for each group are detailed in figure legends). Data were analyzed by t-test, RM one-way ANOVA, two-way ANOVAs and RM two-way ANOVAs, depending on the experimental design, using GraphPad Prism 8. As the electrophysiological recordings were long, some values were missing due to artefacts (3% of the data recorded in extracellular electrophysiology). In this case, data were analyzed by fitting a mixed model proposed by the statistical software. This mixed model uses a compound symmetry covariance matrix, and is fit using Restricted Maximum Likelihood (REML). When indicated, post hoc analyses were carried out with the Bonferroni’s correction procedure. Significance for p values was set at α = 0.05. Effect sizes for the ANOVAs were also reported using partial η^2^ values (Levine and Hullett, 2002; Magnard et al., 2018). Determining these values from the mixed-model analysis was however not accessible.

## Author contributions

RG and SC designed research, RG performed experiments, VC, CC and SC supervised research, RG and SC analyzed data and wrote the manuscript with the help of the other authors.

## Acknowledgments

The authors would like to thank Sabrina Boulet and Yvan Vachez for critical reading of the manuscript. The authors also would like to thank Jacques Brocard and the PIC GIN Platform for technical assistance in fluorescence microscopy and analysis, as well as the in vivo experimental platform.

## Conflict of interest

Authors report no conflict of interest.

## Funding sources

This work was supported by the Institut National de la Santé et de la Recherche Médicale (Inserm), the Agence Nationale de la Recherche (ANR-16-CE16-0002, to SC) and Grenoble Alpes University.

---

**Figure S1.**
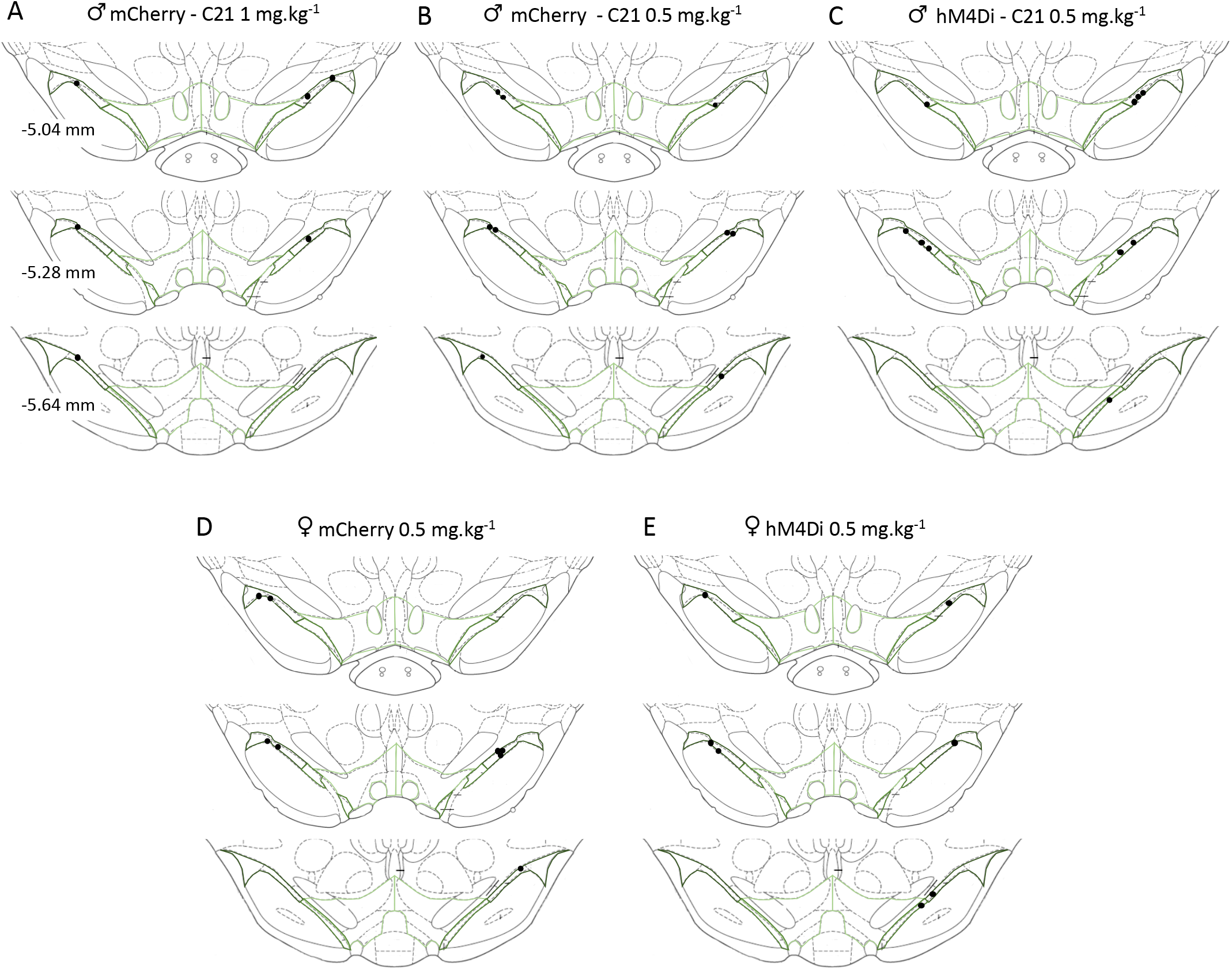
Location of recording sites within the SNc among the different groups studied. Male expressing mCherry treated with 1 mg.kg^−1^ of C21 **(A)** or 0.5 mg.kg^−1^ of C21 **(B)**. Male expressing hM4Di treated with 0.5 mg.kg^−1^ of C21 **(C).** Female expressing mCherry **(D)** or hM4Di **(E)** and treated with 0.5 mg.kg^−1^ of C21.

## Notes

### Competing Interest Statement

The authors have declared no competing interest.

